# Polygene control and trait dominance in death-feigning syndrome in the red flour beetle *Tribolium castaneum*

**DOI:** 10.1101/2021.05.13.443963

**Authors:** Kentarou Matsumura, Takahisa Miyatake

**Affiliations:** Graduate School of Agriculture, Kagawa University, Kagawa, Japan; Graduate School of Environmental and Life Science, Okayama University, Okayama, Japan

**Keywords:** Quantitative trait, death feigning, moving activity, artificial selection, *Tribolium castaneum*

## Abstract

Death-feigning behavior is an anti-predator behavior in a wide range of animal taxa, and it often correlates with the movement (i.e. death-feigning syndrome). In the present study, we performed reciprocal crossing among strains with genetically longer (L strain) and shorter (S strain) duration of death feigning, and investigated related heritable factors in the F_1_ and F_2_ populations. We also investigated moving activity which negatively responded to artificial selection for death feigning in *T. castaneum*. Our results showed that death feigning occurred more frequently and for shorter periods in the F_1_ population. In the F_2_ population, death feigning and movement showed continuous segregation. The distribution of each trait value in the F_2_ generation was different from the distribution of trait values in the parental generation, and no individuals transgressing the distribution of trait values in the parental generation emerged in the F_2_ generation. Chi-square analysis of the observed death feigning and movement of F_1_ and F_2_ progenies rejected the hypothesis of mono-major gene inheritance. These results suggest that death-feigning syndrome is controlled in a polygenic manner. Our study indicated that reciprocal crossing experiments are useful in assessing the quantitative inheritance of behavioral traits.

## 1. Introduction

Animal behaviors are often studied as quantitative traits (Boake 1994; Lynch and Walsh 1998). Reciprocal crossing experiment is a suitable method for studying quantitative traits. When the frequency distribution at F_2_ generation after reciprocal crossing between strains with different trait values does not overlap from the parental generation, it is suggested that the trait is controlled in a polygenic manner. Each gene determines properties that are governed by quantitative inheritance (Falconer and Mackay 1996). If the distribution of the F_2_ population overlaps that of the parent distribution, the trait is likely controlled by a major gene. Furthermore, observation of the distribution at the F1 generation after reciprocal crossing among different strains can estimate which gene is dominant. Therefore, reciprocal crossing can also be used to assess trait dominance that is the phenomenon of one allele of a gene on a chromosome masking or overriding the effect of a different allele of the same gene on the other copy of the chromosome (Falconer and Mackay 1996; Lynch and Walsh 1998). Although the reciprocal crossing method is difficult to reveal the details of the molecular level as RNA-Seq, it is able to investigate the gene inheritance by confirming the actual expression of the traits.

Some previous studies have conducted reciprocal crossing experiment. For example, in cowpea *Vigna unguiculata* (Walp), reciprocal crossing experiments revealed resistance to *Callosobruchus maculatus* (Fabricius) may be inherited as a major gene effects (Redden et al. 1983). In lesser grain borer *Rhyzopertha dominica* (Fabricius), reciprocal crossing with susceptible and resistant lines to phosphine suggested that resistance is no single gene inheritance (Collins et al. 2002). In *Drosophila melanogaster* (Meigen), larval foraging behavior (rover or sitter phenotypes) is controlled by a foraging gene, and reciprocal crossing between rover and sitter lines showed that complete dominance of the rover phenotype in larval behavior (de Belle and Sokolowski 1987). Furthermore, reciprocal crossing of strains with genetically longer or shorter development periods has been conducted in the melon fly *Zeugodacus cucurbitae* (Coquillett); the results showed that shorter developmental period was dominant in the F_1_ population, and that segregation of developmental period in F_2_ did not overlap that of the parent distribution (Miyatake 1997). These results suggest that the developmental period of *Z*. *cucurbitae* is controlled in a polygenic manner (Miyatake 1997). In another study of *Z*. *cucurbitae*, circadian rhythm, which is genetically correlated with developmental period, was also investigated through reciprocal crossing; segregation of the F_2_ population was found to overlap that of the parent generation, suggesting that circadian rhythm is controlled by a major gene in this species (Shimizu et al. 1997). These studies contributed to the discovery of a clock gene in *Z*. *cucurbitae* (Fuchikawa et al. 2010). Therefore, although the reciprocal crossing test is a classic and simple experimental method, it is an important experimental method for investigating quantitative traits.

Death feigning, also known as thanatosis or tonic immobility, is observed in a wide range of animal taxa and is considered to be an adaptive anti-predator behavior (e.g. Edmunds 1974; Miyatake et al. 2009; Humphreys and Ruxton, 2018; Ruxton et al. 2018). A previous study conducted artificial selection for duration of death feigning in the red flour beetle *Tribolium castaneum* (Herbst) and established strains with genetically higher frequency and longer duration of death feigning (i.e., L-strains) and strains with genetically lower frequency and shorter duration of death feigning (i.e., S-strains) (Miyatake et al. 2004). Following encounters with a jumping spider *Hasarius adansoni* (Audouin) as a model predator, L-strain beetles had higher survival rates than S-strain beetles (Miyatake et al. 2004). These findings suggest that death-feigning behavior is an adaptive anti-predator strategy with a genetic basis. Furthermore, L-strain beetles also show less movement than S-strain beetles, suggesting a negative genetic correlation between the duration of death feigning and movement in *T*. *castaneum*; this relationship has been termed the death-feigning syndrome (Miyatake et al. 2008; Matsumura et al. 2017). Other studies have suggested that this genetic behavioral correlation may be controlled by dopamine in *T. castaneum* (Miyatake et al. 2008; Nishi et al. 2010). The death-feigning syndrome has also been observed in *T*. *confusum* (Jaquelin Du Val) (Nakayama et al. 2012; Matsumura et al. 2017) and the adzuki bean beetle *Callosobruchus chinensis* (Linnaeus) (Ohno and Miyatake 2007; Nakayama and Miyatake 2010).

Although L and S strains of death feigning behavior have been established in *T*. *castaneum* by artificial selection (Miyatake et al. 2004; Miyatake et al. 2008), no study has conducted reciprocal crossing experiments to investigate both frequency and duration of death feigning and movement in F_1_ and F_2_ populations. Therefore, in the present study, we conducted reciprocal crossing of L and S strains to investigate segregated phenotypes of death-feigning duration and frequency, as well as movement in F_1_ and F_2_ *T*. *castaneum* populations. The current study provides more data on inheritance of death-feigning behavior, for example, how many genes control this behavior, and how this behavior inherits through generations in field?

## 2. Materials and Methods

### 2.1. Insects and artificial selection for death-feigning duration

In this study, we used previously established L- and S-strain *T*. *castaneum* beetles that had been artificially selected for more than 40 generations (Miyatake et al. 2004; Matsumura and Miyatake 2018). Two replicate lines for each selection regime were created in each strain (i.e. L1, L2, S1, and S2). Detailed methods of the artificial selection and culture of beetles were previously described (Miyatake et al. 2004; Matsumura and Miyatake 2018).

### 2.2. Crossing experiment

We used beetles aged 21-35 days after eclosion in the following experiments. We randomly collected 10 males (virgin) and females (virgin) from each strain (parental generation [P0]) and placed them in a cylindrical plastic container (50 mm in diameter, 30 mm in height) with food until crossing strains (i.e. L1♂ × S1♀, S1♂ × L1♀, L2♂ × S2♀, S2♂ × L2♀) were obtained. In this experiment, the method of mass mating (i.e., 10 males and 10 females) in each strain was adopted instead of individual pairing. Emerging pupae (F_1_) were sexed and placed on Petri dishes (100 mm in diameter, 10 mm in height) according to sex. We randomly collected 10 males (virgin, 21-35 days old) and 10 females (virgin, 21-35 days old) from each cross population (i.e. LS1♂ × LS1♀, SL1♂ × SL1♀, LS2♂ × LS2♀, and SL2♂ × SL2♀) and placed them in a cylindrical plastic container with food. Emerging pupae (F_2_) were sexed and placed on Petri dishes according to sex.

### 2.3. Behavior measurements

We measured the duration and frequency of death feigning and movement in each population (P_0_, F_1_, and F_2_). Virgin males (21-35 days old) and females (21-35 days old) were randomly collected from each population and placed in a 48-well tissue plate with food. The next day, a beetle was gently moved to a dish (110 mm in diameter) and touched with a wooden stick. When the beetle shows tonic immobility by the artificial stimuli, we defined of immobility as death feigning behavior and measured duration of immobility. If the beetle did not show death feigning, the stimulus was repeated for two times. When the beetle did not show death feigning even by total these stimuli, we recorded “no death feigning” beetle (see also Miyatake et al. 2004). Observations of death feigning were conducted during 12:00 and 18:00 in the room maintained 25°C.

To measure movement of *T. castaneum*, we used an image tracking system (DIGIMO, Japan). The day before this experiment, virgin males and females were randomly corrected from each population and placed in a 48-well tissue plate with food. Each beetle was gently moved to a Petri dish (30 mm in diameter) 2h before the measurement. Movie of walking movement was recorded using a video CCD camera (SEYE130SN, Science-eye, Japan) for 30 min. Moving distance was analysed by software (2D-PTV Ver. 9.0, Digimo, Japan), and we defined this moving distance as movement. All measurements were conducted during 10:00 and 18:00 at room maintained 25°C (see also Matsumura and Miyatake 2015).

### 2.4. Statistical analyses

Data for each generation were analysed separately. The death-feigning duration (s) and movement (mm) were analysed using a generalised linear mixed model (GLMM) with gamma distribution, strain as a fixed effect, and replicate line as a random effect. Death-feigning frequency was analysed using a GLMM with binomial distribution.

Observed data were tested against expected data using chi-square test for goodness-of-fit. Expected values were calculated from classical Mendelian inheritance that assuming monogenic inheritance with complete dominance (i.e., 1:2:1 or 3:1 in F_2_ generation). This classical analysis method was used in many previous studies (see Harris 1912; Snedecor and Cochran 1967). In death-feigning duration and walking distance, a frequency distribution of 1:2:1 is assumed in the F_2_ generation as the mono-gene inheritance (Fig. S1, S3). In death- feigning frequency, a frequency distribution of 3:1 is assumed in the F_2_ generation as the mono-gene inheritance for the threshold trait (Fig. S2). To compare of distributions observed and null hypothesis, we used chi-square goodness of fit test (see Snedecor and Cochran 1967). All analyses were conducted using R ver. 3.4.3 software (R Core Team 2017).

## 3. Results

The distributions of death-feigning duration in the P_0_, F_1_, and F_2_ populations are shown in Fig. 1. In the P_0_ populations, death-feigning behavior was significantly longer in L-strain beetles than in S-strain beetles (Fig. 1, Table 1). The duration of death-feigning behavior differed significantly between the F_1_ reciprocal crosses (L♂ × S♀ vs. S♂ × L♀; Table 1 and Fig. 1). Because the mean death-feigning duration of S-strain beetles was closer to the mean of the F_1_ population than that of L-strain beetles (Table S1), shorter death-feigning duration appears to be partially dominant. F_2_ population showed a similar distribution to that of the F_1_ population that showed continuous segregation of the death-feigning duration trait (Fig. 1). Because the distribution of death-feigning duration in the F_2_ population did not overlap those of the P_0_ populations, death-feigning duration is likely not controlled by a major gene. Together, these segregation patterns suggest that death-feigning duration is controlled in a polygenic manner in *T*. *castaneum*. We detected no significant differences between crossed F_2_ populations (Table 1).

**Figure 1.**
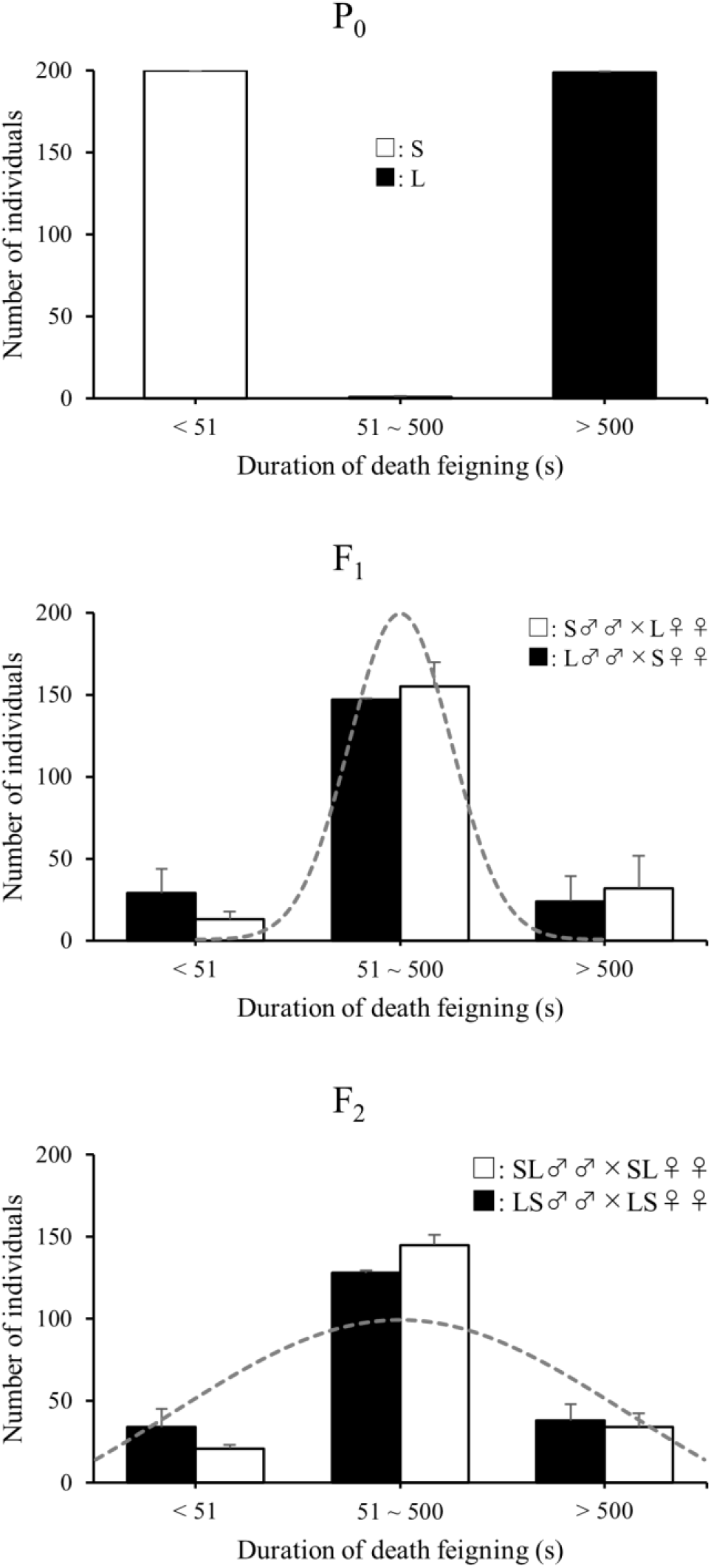
Distributions of duration of death feigning at P_0_, F_1_, and F_2_ generations reciprocal crossing experiments. Error bars showed standard error. The grey curves showed the theoretical distribution assuming quantitative inheritance.

**Table 1.** Results of GLMM (strain as a fixed effect, replicate line as a random effect) for duration and frequency of death feigning and moving distance of beetles from each generation.

| Behavior | Generation | *d.f.* | $\chi^2$ | *p* |
| --- | --- | --- | --- | --- |
| Death-feigning duration | P<sub>0</sub> | 1 | 15358.00 | < **0.0001** |
|  | F<sub>1</sub> | 1 | 18.55 | < **0.0001** |
|  | F<sub>2</sub> | 1 | 1.01 | 0.3144 |
| Death-feigning frequency | P<sub>0</sub> | 1 | 291.69 | < **0.0001** |
|  | F<sub>1</sub> | 1 | 0.34 | 0.5585 |
|  | F<sub>2</sub> | 1 | 2.13 | 0.1442 |
| Moving activity | P<sub>0</sub> | 1 | 154.30 | < **0.0001** |
|  | F<sub>1</sub> | 1 | 1.00 | 0.3168 |
|  | F<sub>2</sub> | 1 | 0.45 | 0.5041 |

The death-feigning frequency distribution of each population is shown in Fig. 2. In the P_0_ populations, L-strain beetles had a significantly higher frequency of death-feigning behavior than S-strain beetles (Fig. 2, Table 1). Among the F_1_ populations, the vast majority of individuals feigned death (Fig. 2). F_2_ populations also displayed a higher frequency of death-feigning behavior (Fig. 2, Table S1).

The distribution of movement measurements in each population are shown in Fig. 3. Movement measurement frequency was less densely distributed than death-feigning frequency. In the P_0_ populations, S-strain beetles moved significantly more frequently than L-strain beetles (Fig. 3, Tables 1). In F_1_ populations, we detected no significant differences in movement measurement frequency among reciprocal crosses (Fig. 3, Table S1). Continuous segregation of movement behavior was observed in F_2_ populations and did not overlap with that of the P_0_ populations (Fig. 3). These segregation patterns suggest that movement is controlled in a polygenic manner in *T*. *castaneum*. No significant differences in movement behavior were detected among F_1_ and F_2_ crossed populations (Table 1).

Table 2 showed results of chi-square analysis for test of differences between observed data and theoretical model of monogenic inheritance in each behavioral traits. Observed data of death feigning and moving showed similarly distributions with theoretical predicted distributions (1:2:1 or 3:1) in F_1_ and F_2_ generation, respectively (grey curves in Fig. 1-3). Chi-square analysis of the observed data rejected the hypothesis of mono-major gene inheritance in duration of death feigning (Fig. 1, Table 2) and moving distance (Fig. 3, Table 2). In frequency of death feigning, chi-square analysis did not reject the hypothesis of mono-major gene inheritance at F_1_, but rejected at F_2_ generation (Fig. 2, Table 2).

**Table 2.**
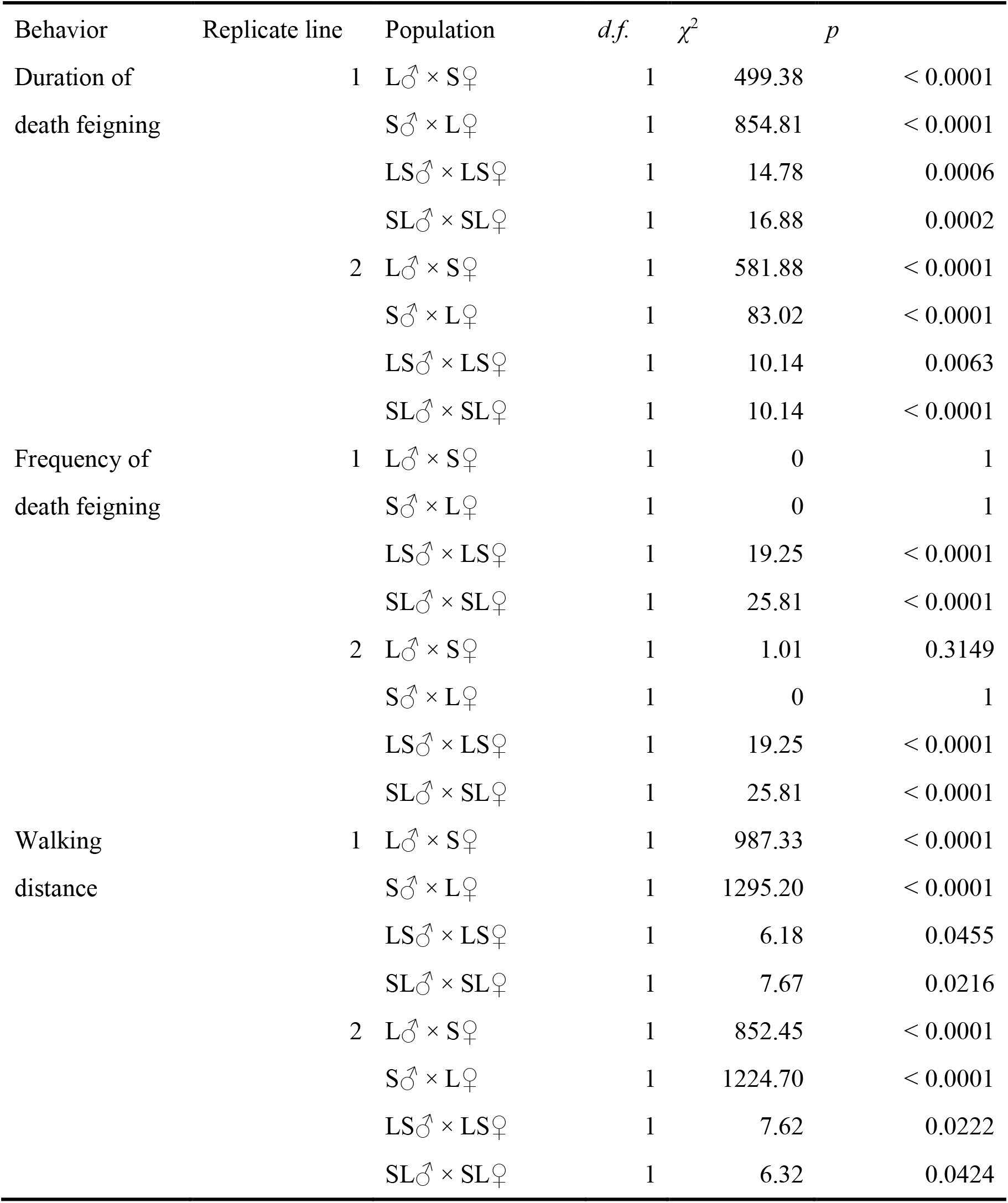
Results of chi-square analysis for tests of differences between observation value and null hypothesis in duration and frequency of death feigning and walking distance, respectively.

## 4. Discussion

The reciprocal crossing of L- and S-strain *T*. *castaneum* beetles showed that F_1_ populations were similar to S-strain beetles in terms of shorter death-feigning duration (Fig. 1, Table S1) and higher frequency (Fig. 2, Table S1). Therefore, these results suggested that shorter duration and higher frequency of was dominant in death-feigning behavior. On the other hand, movement behavior in each reciprocal crossing in the F_1_ populations showed no particular trend (Fig. 3). Thus, reciprocal crossing showed different results among the behavioral traits examined in this study in the F_1_ populations.

Previous studies have conducted reciprocal crossing experiments to explore the inheritance of behavioral traits. For example, in the melon fly *Z. cucurbitae*, crossing strains selected for longer or shorter development periods showed that shorter developmental period was dominant in F_1_ populations (Miyatake 1997). In the cowpea weevil *C. maculatus*, reciprocal crossing of strains selected for different dispersal ability revealed no dominance in F_1_ populations (Messina 1987). In the field vole *Microtus agrestis* (Linnaeus), reciprocal crossing of populations with higher and lower activity showed no dominance in activity among F_1_ populations (Rasmuson et al. 1977). Thus, whether there is dominant in a behavioral trait may vary across its behaviors and species.

Our results suggest that death-feigning behavior is genetically more stable with shorter duration and higher frequency. However, when the beetle encounters with predators, beetles with longer death-feigning duration tend to have higher survival rates than those with shorter death-feigning duration. Indeed, the beetles from a population coexist with predators showed higher frequency and longer duration of death-feigning behavior in *T. castaneum* (Konishi et al. 2020). However, higher frequency and longer duration of death feigning has physiological costs. Kiyotake et al. (2013) found that L-strain beetles were more sensitive to thermal and vibrational stress than S-strain beetles. Indeed, beetles from laboratory populations without predators showed shorter duration and lower frequency of death feigning. These results suggest that frequency and duration of death-feigning behavior is shaped by ecological and physiological factors as well as genetic factors.

In F_2_ populations, continuous segregation of death-feigning duration did not overlap with that of the P_0_ populations (Fig. 1), suggesting that duration is controlled in a polygenic manner. Because movement measurement frequency showed continuous segregation in F_2_ reciprocal crossing, and the F_2_ distributions did not overlap with the P_0_ distributions, movement is also controlled in a polygenic manner in *T*. *castaneum*. Furthermore, chi-square tests showed that death-feigning behavior and movement are not controlled by mono-major gene (Fig. 1-3, Table 2). In *D. melanogaster*, chi-square analysis was conducted to test the differences between observed ratios by reciprocal crossing between rover and sitter lines and expected Mendelian rations. Because observed ratios were not significantly different from expected Mendelian ratios in all reciprocal cross, it is suggested that the larval foraging behavior was controlled by a single gene (de Belle and Sokolowski 1987). In a previous study investigating the inheritance of three isozymes in peanut, the segregation ratios of F_2_ from crosses between different lines were compared with the expected ratios by chi-square analysis, and it was found that some isozymes showed Mendelian inheritance and others are not (Grieshammer and Wynne 1990). In the melon fly *Z. cucurbitae*, because frequency distribution of F_2_ population after reciprocal crossing between longer- and shorter-developmental period strains did not overlap with parent distribution, it was suggested that the developmental period of *Z. cucurbitae* is controlled in a polygenic manner (Miyatake 1997). In another study of *Z. cucurbitae*, because segregation of circadian rhythm of the F_2_ population after reciprocal crossing was overlapped with the parent generation, it was suggested that the circadian rhythm is controlled by a major gene (Shimizu et al. 1997). These results indicated that the reciprocal crossing test is a suitable experimental method for investigating quantitative traits.

In the present study, we did not conduct back-crossing experiments. Because our results suggested that the L phenotype requires homozygous locus (or loci), back-crossings could have provided more information with larger phenotypic variations. We need additional study that conducts of back-crossing experiments using the L- and S-strains in the future. Moreover, we conducted reciprocal crossing experiment by mass mating using 10 individuals of each sex but not single-pair mating. Additional studies are needed that single pair mating with backcrossing to provide a high-resolution phenotype data offering more inferences in the genetics. We did not assess the repeatability (i.e., intra-individual variation) of death-feigning behavior. The data of repeatability combined with results of this study may provide more useful information. It will be necessary to carry out these experiments in the future.

Miyatake et al. (2008) reported that in *T*. *castaneum*, L-strain beetles showed significantly lower brain dopamine expression than S-strain beetles. Injection of dopamine into L-strain beetles decreased their death-feigning duration (Nishi et al. 2010). These previous findings suggest that death-feigning behavior is controlled by dopamine-related genes. A recent study conducted transcriptome analysis of gene expression in L- and S-strain *T*. *castaneum* beetles, and found significant differences between strains in insulin- and longevity-related genes, as well as dopamine-related genes (Uchiyama et al. 2019). These findings and those of the present study suggest that death-feigning behavior is controlled by multiple genes including physiological trait-related and longevity-related genes. Our results are important not only because they support the previous results suggested by its gene expression analysis, but also reveal the dominance of death-feigning and movement and that these behaviors are not controlled by major gene.

Death-feigning syndrome is observed in many species including the Japanese quail *Coturnix japonica* (Temminck and Schlegel) (e.g., Mills and Faure 1991; Recoquillay et al. 2013) and guinea pig *Cavia porcellus* (Linnaeus) (e.g., Thompson et al. 1981; Leite-Panissi et al. 2006), as well as insects (e.g., Ohno and Miyatake 2007; Nakayama et al. 2010). Although death-feigning syndrome has been observed throughout a wide range of animal taxa, the present study is the first study that conducted reciprocal crossing experiment. Further study is required to investigate the mode of inheritance of death-feigning syndrome.

## Acknowledgement

This work was supported by Grant-in-Aid for Japan Society for the Promotion of Science (JSPS) Fellows 20J00383 to K.M. and 18H02510 to T.M.

## Compliance with ethical standards

### Ethical approvals

The laboratory population of *T. castaneum* used in this study have maintained at Okayama University. This population has been maintained on whole meal flour with yeast. We reared this population at 25 °C, which resemble natural conditions for this insect. All animals in the study were handled more carefully. The use of these animals conforms to the Animal Ethics Policy of Okayama University.

## Supplementary material

**Figure S2.**
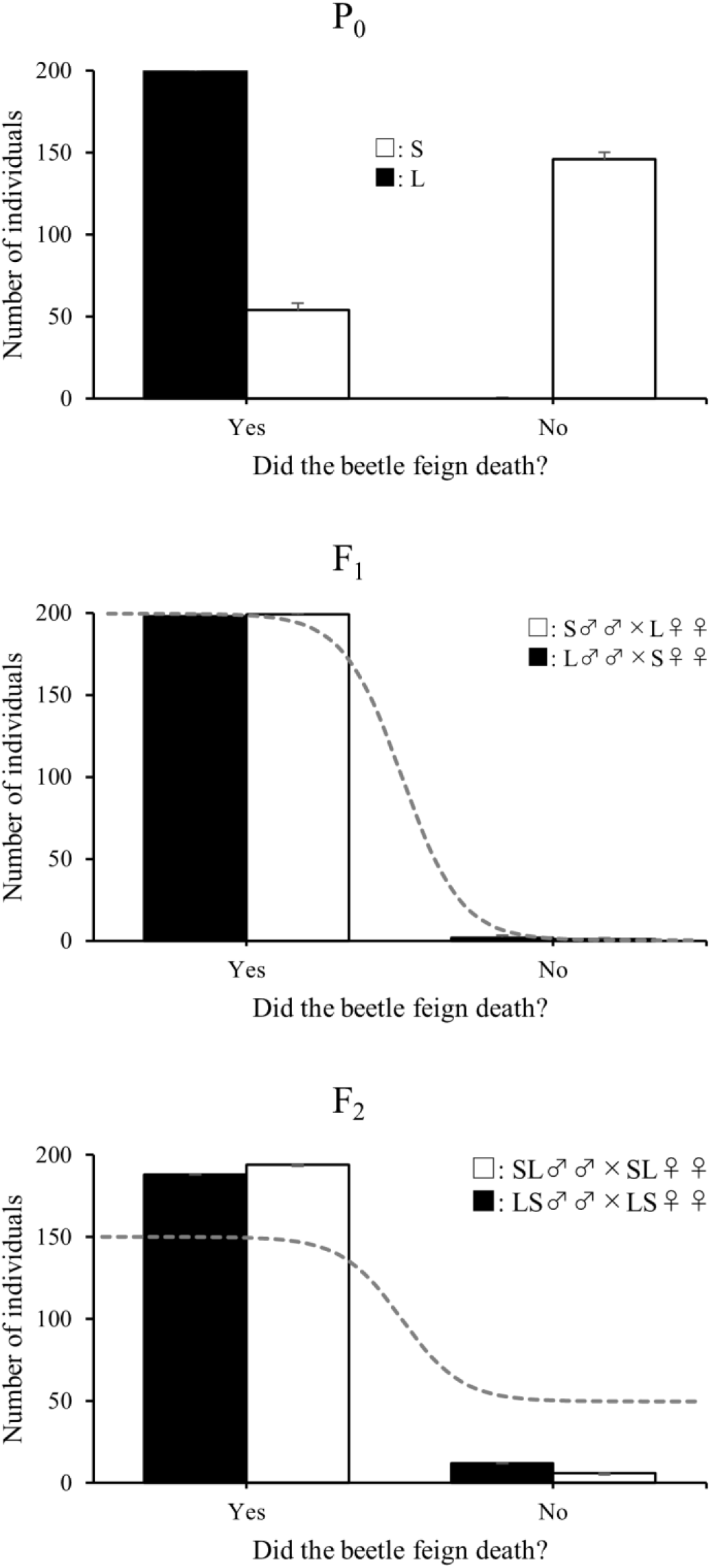
Distributions of number of individuals who feign death or not duration of death feigning at P_0_, F_1_, and F_2_ generations reciprocal crossing experiments. Error bars showed standard error. The grey curves showed the theoretical distribution assuming quantitative inheritance.

**Figure S3.**
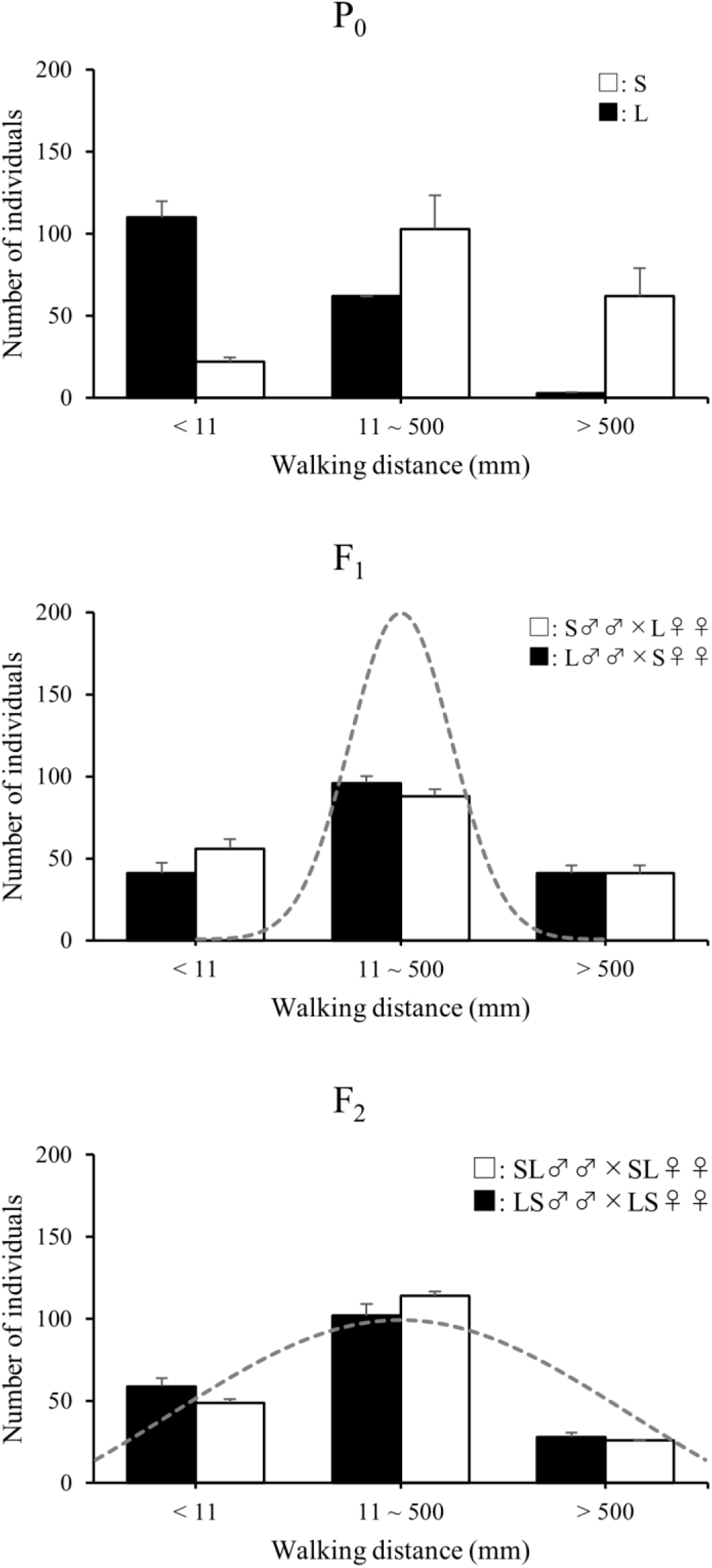
Distributions of walking distance duration of death feigning at P_0_, F_1_, and F_2_ generations reciprocal crossing experiments. Error bars showed standard error. The grey curves showed the theoretical distribution assuming quantitative inheritance.

**Table S1.** Mean ± SD of each behavioral trait from each population at P_0_, F_1_, and F_2_ generation, respectively. Each value of population was contained sex.

| Replicate line | Generation | Population | Mean ± SD | | |
| --- | --- | --- | --- | --- | --- |
|  |  |  | DF duration | DF frequency | Moving distance |
| 1 | P<sub>0</sub> | L | 6769.30 ± 4211.84 (100) | 1.00 (100) | 53.06 ± 123.74 (80) |
| | | S | 0.60 ± 1.76 (100) | 0.30 (100) | 293.91 ± 302.93 (94) |
| | F<sub>1</sub> | L♂ × S♀ | 363.25 ± 279.72 (100) | 1.00 (100) | 237.18 ± 325.62 (87) |
| | | S♂ × L♀ | 501.24 ± 673.27 (100) | 0.99 (100) | 208.042 ± 320.99 (96) |
| | F<sub>2</sub> | LS♂ × LS♀ | 486.54 ± 953.33 (100) | 0.94 (100) | 208.45 ± 313.93 (95) |
| | | SL♂ × SL♀ | 334.93 ± 530.29 (100) | 0.97 (100) | 175.22 ± 252.58 (95) |
| 2 | P<sub>0</sub> | L | 6343.50 ± 3805.72 | 1.00 (100) | 59.06 ± 142.90 (95) |
| | | S | 0.66 ± 1.87 | 0.24 (100) | 575.73 ± 652.81 (93) |
| | F<sub>1</sub> | L♂ × S♀ | 116.31 ± 92.55 | 0.98 (100) | 316.87 ± 338.72 (91) |
| | | S♂ × L♀ | 170.46 ± 157.04 | 1.00 (100) | 269.02 ± 307.25 (89) |
| | F<sub>2</sub> | LS♂ × LS♀ | 187.87 ± 209.90 | 0.94 (100) | 222.61 ± 405.68 (94) |
| | | SL♂ × SL♀ | 219.44 ± 264.40 | 0.97 (100) | 212.81 ± 296.69 (94) |

